# RevelioPlots: An Interactive Web Application for Fast AI-Based Protein Models Quality Assessment

**DOI:** 10.64898/2026.02.23.707401

**Authors:** Leonardo Luiz de Sena Fernandes, Arthur Henrique Dantas de Azevedo, João Vitor Santos de França, João Paulo Matos Santos Lima

**Affiliations:** Bioinformatics Multidisciplinary Environment (BioME), Digital Metropolis Institute (IMD), Universidade Federal do Rio Grande do Norte (UFRN), Natal, RN, 59078-900, Brazil

**Keywords:** protein structure, model evaluation, Ramachandran’s Plot, stereochemical quality, mmcif files

## Abstract

High-accuracy protein structure prediction by deep learning requires rigorous model quality assessment, a process currently hampered by fragmented, non-interactive tools designed for older experimental data formats. We present RevelioPlots, an open-source, interactive web application (Python/Streamlit) that simplifies and streamlines the assessment of AI-predicted protein structure quality. Its key feature is the combination of statistical pLDDT score analysis (mean, median, box plots) with an interactive, confidence-colored Ramachandran plot. This integration establishes a direct visual link between a model’s predicted local reliability (pLDDT) and its stereochemical feasibility (backbone geometry). RevelioPlots handles both individual and batch-uploaded models, intelligently falling back to B-factors as a proxy for pLDDT values. Using example model proteins, we demonstrated the tool’s effectiveness, revealing differences in reliability and a clear visual correlation between regions of low pLDDT scores and residues in sterically disallowed regions. By unifying these critical metrics, RevelioPlots empowers non-experienced researchers to quickly and intuitively assess, compare, and interpret structural model quality, enabling a more confident and integrated use of predicted data.

**Availability:** RevelioPlots is available at revelioplots.streamlit.app, with the source code publicly accessible on GitHub at https://github.com/evomol-lab/RevelioPlots.

## 1 BACKGROUND AND MOTIVATION

High-accuracy protein structure prediction has been revolutionized by deep learning methods such as AlphaFold (Jumper et al. 2021), RoseTTAFold (Baek et al. 2021; 2023), and Boltz (Wholwend et al. 2024; Passaro et al. 2025). However, unlike experimental structures determined via physical measurement, AI-generated models are *in silico* hypotheses. This distinction makes rigorous quality assessment a non-negotiable step before their use in downstream applications.

This evaluation is a complex process relying on distinct yet complementary approaches. It begins by assessing the model’s reliability using per-residue confidence metrics, such as the predicted Local Distance Difference Test (pLDDT) scores (Jumper et al. 2021). Based on lDDT-C⍰, this assessment then broadens to global quality scores, which are determined by comparing the model’s statistical potentials and distance constraints against databases of known experimental structures (Mariani et al. 2013). Crucially, the model must also pass a rigorous stereochemical validation to confirm its biological feasibility. This step uses tools such as Ramachandran plots to verify that the model adheres to the fundamental principles of molecular geometry (e.g., valid bond angles and steric non-repulsion) (Ramachandran et al., 1963).

The current challenge is the highly fragmented validation workflow. To perform a comprehensive quality assessment, researchers typically employ a wide array of tools, such as SWISS-MODEL’s quality assessment module (Tunyasuvunakool et al. 2021), Procheck (Laskowski et al. 1993), What_Check (Hooft et al. 1996), Verify3D (Lüthy et al. 1992), ERRAT (Colovos et al. 1993), and Molprobity (Williams et al. 2018). Despite their utility, these platforms generally lack interactive visualizations that enable the quick identification of problematic regions and a direct correlation with stereochemical quality. Additionally, many of these tools were designed for experimental data, expecting PDB files with B-factor values instead of modern formats with embedded confidence scores. This forces researchers to navigate a cumbersome process of file conversions and disconnected analyses, hindering rapid and integrated assessment.

RevelioPlots addresses this gap by providing a simple and user-friendly interactive web application for intuitive visualization, analysis, and comparison of protein structure models. By focusing on the direct correlation between pLDDT scores and backbone conformation, RevelioPlots streamlines the entire quality assessment pipeline. In addition, the tool targets non-experienced users who are taking advantage of the high accessibility of AI-based protein structural modelling methods.

## 2 REVELIOPLOTS IMPLEMENTATION

### Architecture and Technology

RevelioPlots is an open-source web application developed in Python 3.8+ and built upon the Streamlit framework for its interactive interface (Streamlit Inc., 2025). Core functionalities are handled by established bioinformatics and data science libraries, including Biopython (Cock et al. 2009), Pandas (McKinney, 2010), NumPy (Harris et al. 2020), and Plotly (Plotly Technologies Inc. 2015). The application is available at revelioplots.streamlit.app, and can also be executed locally, ensuring data privacy and responsive performance.

### Input Data Processing

RevelioPlots supports both the analysis of individual structures and the comparative assessment of multiple models uploaded as a batch. All models must be provided in the Crystallographic Information File (.cif or.mmcif) format. Once uploaded, it parses these files and extracts essential information for each residue: its identity (e.g., ALA, GLY), atomic coordinates for calculating backbone dihedral angles (Φ and Ψ), and the associated confidence score.

A key feature of this pipeline is its flexible handling of confidence metrics. The application first searches for explicit pLDDT scores. If these are not present, it intelligently falls back to using the B-factor column as a proxy. This interpretation is based on the strong conceptual correlation between these metrics: high B-factors in experimental structures typically denote regions of high flexibility or disorder, which are the same regions that deep learning models typically predict with low confidence (low pLDDT). Therefore, the tool assumes an inverse relationship between B-factor values and the inferred confidence score (Afonine et al., 2023).

### Features & Visualization

#### pLDDT Score Analysis

Provides a statistical summary (mean, median, standard deviation) and an interactive box plot showing the distribution of pLDDT scores across all residues. pLDDT is a per-atom or per-residue confidence estimate, ranging from 0 to 100, where higher values indicate greater confidence in the local predicted structure. This provides a rapid, high-level overview of the model’s overall quality, allowing researchers to quickly identify disordered or poorly modeled regions and check if the prediction is generally reliable before proceeding with more detailed analyses.

#### Confidence-Colored Sequence View

Displays the full amino acid sequence where each residue is colored according to its pLDDT score using the standard AlphaFold color scheme (Blue: >90 high confidence; Cyan: 70-90 confident; Yellow: 50-70 low confidence; Red: <50 very low confidence/likely disordered). This view includes a positional scale and interactive tooltips for residue details, which are critical for immediate localization of specific low-confidence residues or domains within the context of the primary sequence, facilitating hypothesis generation about flexible loops or disordered regions.

#### Interactive Ramachandran Plot

Visualizes the Phi (Φ) versus Psi (Ψ) dihedral angles for each residue. Points are colored by their pLDDT value, allowing for direct correlation between local backbone conformation and model confidence. Shaded regions indicate sterically allowed conformations, and interactive tooltips provide detailed residue information that establishes a direct correlation for distinguishing between genuinely strained conformations, which may be functionally important, and geometric artifacts resulting from low model confidence (outliers with low pLDDT scores), except for glycine and proline residues, which are expected to fall outside the main regions.

### Multi-Structure Comparative Analysis

In this mode, the application first presents a comparative overview of pLDDT statistics and distributions for all uploaded models. Following this summary, a “Per-Structure Analysis” section provides collapsible panes for each model, allowing users to perform a detailed inspection of its individual Confidence-Colored Sequence and Ramachandran Plot. This hierarchical approach enables researchers to efficiently move from a high-level comparison to a granular investigation, quickly identifying the most reliable model or pinpointing regions of significant conformational or confidence variance.

## 3 VISUAL AND INTERACTIVE ANALYSIS

To showcase the comparative features of RevelioPlots, we utilized the application’s **built-in example datasets**. These consist of two distinct proteins, each represented by two structural model series: sodc (superoxide dismutase [Cu-Zn] from *Drosophila melanogaster*, Uniprot ID: P61851) and pct (an uncharacterized protein from Trypanosoma equiperdum, GenBank accession XP_067078569) that reflect hypothetical outputs from different deep learning-based prediction algorithms. These pre-loaded models allow users to perform a direct assessment of relative quality and familiarize themselves with the analytical workflow before processing their own data (**Figure 1**).

**Figure 1.**
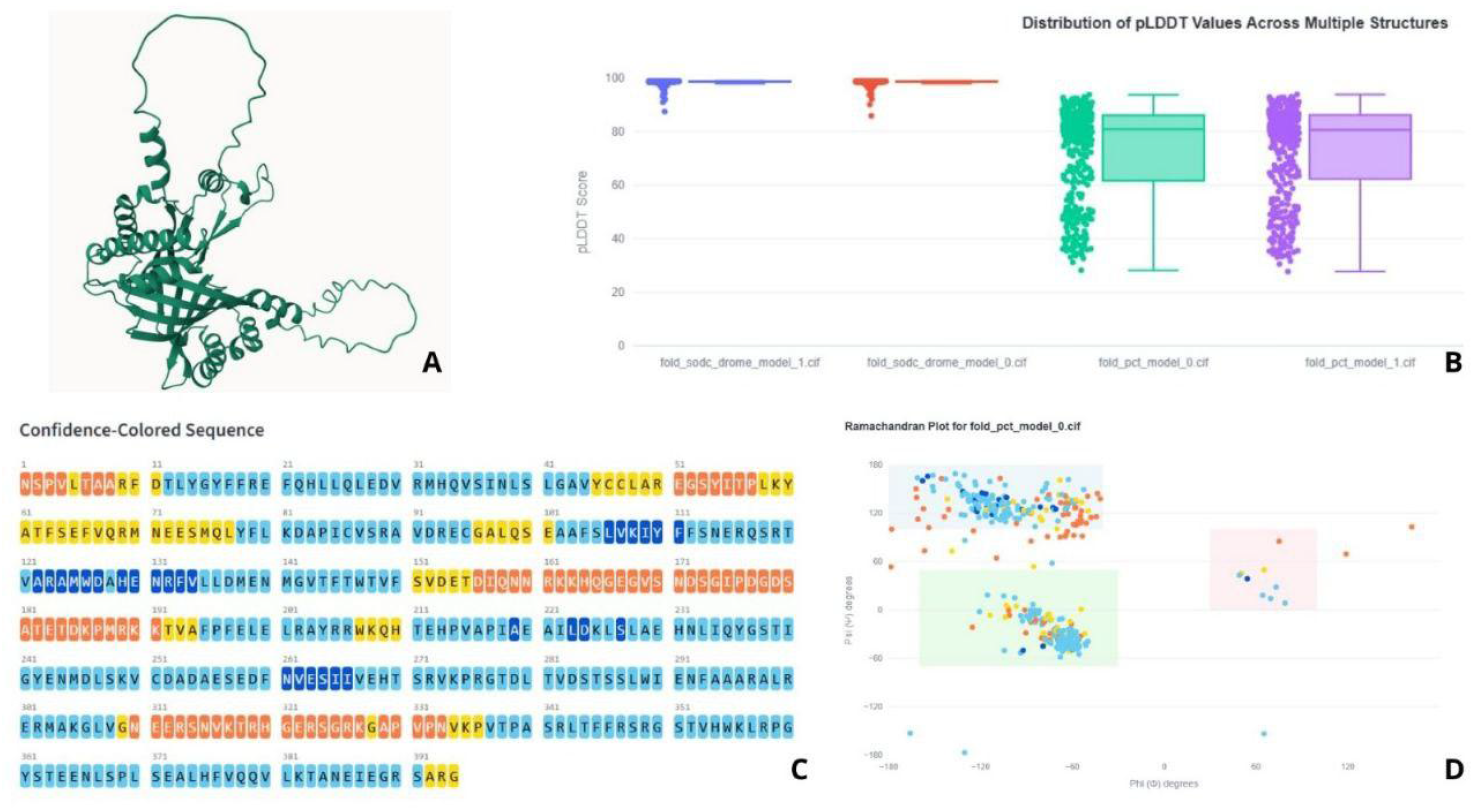
Multi-structure comparison analysis made by RevelioPlots. **(A)** Three-dimensional model of the example protein (pct_model_0) in the tool. **(B)** Comparative statistics boxplot among the different structural models used as examples in the tool, allowing visualization of structural dispersion and variability across the generated models. **(C)** Primary protein sequence colored residue-by-residue according to the calculated confidence interval, emphasizing structurally reliable regions and segments with higher predictive uncertainty. **(D)** Individual residue mapping on the Ramachandran plot, with points colored according to the same confidence interval shown in (C), enabling correlation between conformational quality and structural robustness at the residue level.

The integrated 3D structural visualization **(Fig. 1A)** provides the first spatial context for this analysis. For the **pct-0** model, the viewer reveals a well-folded globular core, but also highlights long, extended terminal segments. This interactive 3D environment allows users to rotate and zoom into the structure, visually mapping the overall topology before deep-diving into per-residue confidence and stereochemical metrics.

The initial statistical analysis allows researchers to compare pLDDT metrics across all pre-loaded models. In this demonstration, the tool revealed a clear distinction in model reliability: the **sodc** models exhibited exceptionally high confidence (mean of ∼98.30), while the pct models showed significantly lower, more variable scores. Specifically, **pct-0** achieved a mean pLDDT of 72.79 and a median of 80.97. The comparative box plot (**Fig. 1B**) visually highlights this discrepancy, showing the pct models’ wide variance and high standard deviation (17.62) compared to the near-perfect scores of the **sodc** group.

The Confidence-Colored Sequence View of the pct-0 model, exemplified in Figure 1C, enables immediate localization of specific low-confidence residues or domains. In this model, the sequence reveals a high-confidence core region colored in blue, which stands in stark contrast to the N-terminal and C-terminal regions. These terminals are colored yellow and red, indicating low to very low confidence (pLDDT < 70), a result consistent with intrinsically disordered regions or flexible loops. This view is critical for generating hypotheses about which parts of the protein should be interpreted with caution.

Finally, the interactive Ramachandran plot (Fig. 1D) provides the definitive stereochemical context for the pct-0 model. Points corresponding to the high-pLDDT core domain occupy the sterically allowed regions, confirming their geometric quality. Conversely, residues from the low-confidence terminal regions are scattered across the plot, with many falling into disallowed regions of conformational space. This direct visual correlation between low pLDDT scores and poor backbone geometry underscores the likelihood that these regions do not maintain a stable fold. This integration allows users to distinguish between genuinely strained conformations and geometric artifacts resulting from low model confidence.

## 4 CONCLUSION

Here, we presented RevelioPlots, an interactive web application designed to simplify and streamline the quality assessment of AI-predicted protein structures. By combining statistical analysis of pLDDT scores with an interactive, confidence-colored Ramachandran plot, our tool provides a direct visual link between model reliability and stereochemical feasibility. RevelioPlots addresses the challenge of a fragmented workflow, empowering non-experienced researchers to quickly and intuitively assess, compare, and interpret the quality of structural models, enabling a more confident use of predicted data in scientific investigations.

## Availability

RevelioPlots can be executed directly at revelioplots.streamlit.app. The tool source code is publicly available on GitHub at https://github.com/evomol-lab/RevelioPlots. The repository includes detailed installation instructions and example files to facilitate testing.

## Acknowledgements

The authors are indebted to the High-Performance Computing Center (NPAD) at UFRN for providing computational resources.

## Funding

This study was funded by the Coordenação de Aperfeiçoamento de Pessoal de Nível Superior - Brasil (CAPES). LLS Fernandes was supported by a CAPES scholarship - Finance Code 001.

## Artificial Intelligence Disclosure

The authors used generative AI technologies for code auditing and performance optimization (Gemini® and GitHub-CoPilot®) and for stylistic editing of the manuscript (Grammarly®). Throughout this process, the authors maintained full control over the research design and interpretation of results; the AI acted solely as a technical and linguistic aid.

## Conflict of Interest

none declared.

## Notes

https://github.com/evomol-lab/RevelioPlots

https://revelioplots.streamlit.app

### Competing Interest Statement

The authors have declared no competing interest.

